# Centromere repositioning induced by inner kinetochore impairment generates a meiosis barrier

**DOI:** 10.1101/401257

**Authors:** Min Lu, Xiangwei He

## Abstract

Centromeres dictate the sites for kinetochore assembly on chromosomes, while their own position on each chromosome is determined epigenetically by a specific histone H3 variant CENP-A. For all eukaryotic species, the chromosomal position of each centromere is distinct and inherited with high fidelity, although the mechanisms underlying the epigenetic stability and its functional significance remain largely unknown. Here in the fission yeast *Schizosaccharomyces pombe*, we show that mutations in inner kinetochore components influence centromeric chromatin organization to various levels. In extreme cases, a single deletion of *wip1, mhf1* and *mhf2* (the conserved CENP-T-W-S-X complex subunits) or double deletions of *cnp3* (a homologue of mammalian CENP-C) and *fta6* (a *pombe* specific component) induce centromere repositioning - inactivation of the original centromere and formation of a neocentromere - in one of the three chromosomes at random. Neocentromeres tend to locate in pericentromeric heterochromatin regions, although heterochromatin is not required for centromere inactivation. Cells carrying a neocentromere are competent in mitosis and in meiosis of homozygotes. However, when these cells are crossed to cells carrying the original centromere, the progeny suffers severe lethality due to defects in meiotic chromosome segregation. These results recapitulate a meiosis barrier that could initiate genetic divergence between two populations with mismatched centromeres, documenting a potential role of the Evolutionary New Centromeres (ENCs) in speciation.

**Significance Statement:** In eukaryotes, centromeres are chromosomal regions where kinetochores are assembled and the positions of centromeres are accurately inherited. While the centromere and kinetochore assembly are extensively studied, the mechanisms that each centromere maintain its identity on chromosomes are still not well understood. In this study, we demonstrated that the inner kinetochore is required for the normal centromere identity as single depletion of the inner kinetochore CENP-T-W-S-X complex or double deletions of *cnp3*/CENP-C and *fta6* induce centromere repositioning. We further showed cells carrying a neocentromere are reproductively isolated from the wildtype population carrying the original centromere. Taken together, these results suggest that induced centromere repositioning mimics the evolutionary new centromeres and is sufficient to cause reproductive isolation.

---

Centromeres are specialized chromosomal regions where kinetochores are assembled (1), their positions are determined epigenetically by a specific histone H3 variant CENP-A. Fission yeast exhibits a characteristic centromeric chromatin organization pattern (2). The central cores, consisting of mostly unique DNA sequences (*cnt*) and part of the innermost repeats (*imr*), are occupied by Cnp1/CENP-A nucleosomes interspersed with canonical H3 nucleosomes; whereas the flanking regions comprise repetitive DNA sequences (outermost repeats *otr* and part of *imr*) are packed into heterochromatin (3, 4), as marked by histone H3 lysine 9 methylation (H3K9me2) (5). The boundaries between the heterochromatin and the central cores are strictly delimited by *tDNA* elements (Fig. 1 A and B, Diagrams) (6). The inner kinetochore is proximal to or in direct contact with the CENP-A nucleosomes, linking centromeres to the outer kinetochore, which in turn physically binds the spindle microtubules (7). Certain inner kinetochore components (*e.g.*, Mis6 and Cnp3 in *S. pombe*; CENP-C and CENP-N in vertebrates) are required for maintaining proper levels of CENP-A nucleosomes in centromeres (8-11). Moreover, partial dysfunction of kinetochore (*e.g., mis6-302, mis12-537* and *ams2*Δ) facilitates centromere inactivation and rescues the high rates of lethality caused by an engineered dicentric chromosome in fission yeast (12). Overall, despite extensive knowledge of centromeric chromatin organization and kinetochore assembly, how centromeres establish and maintain their epigenetic identity remains opaque.

**Fig. 1.**
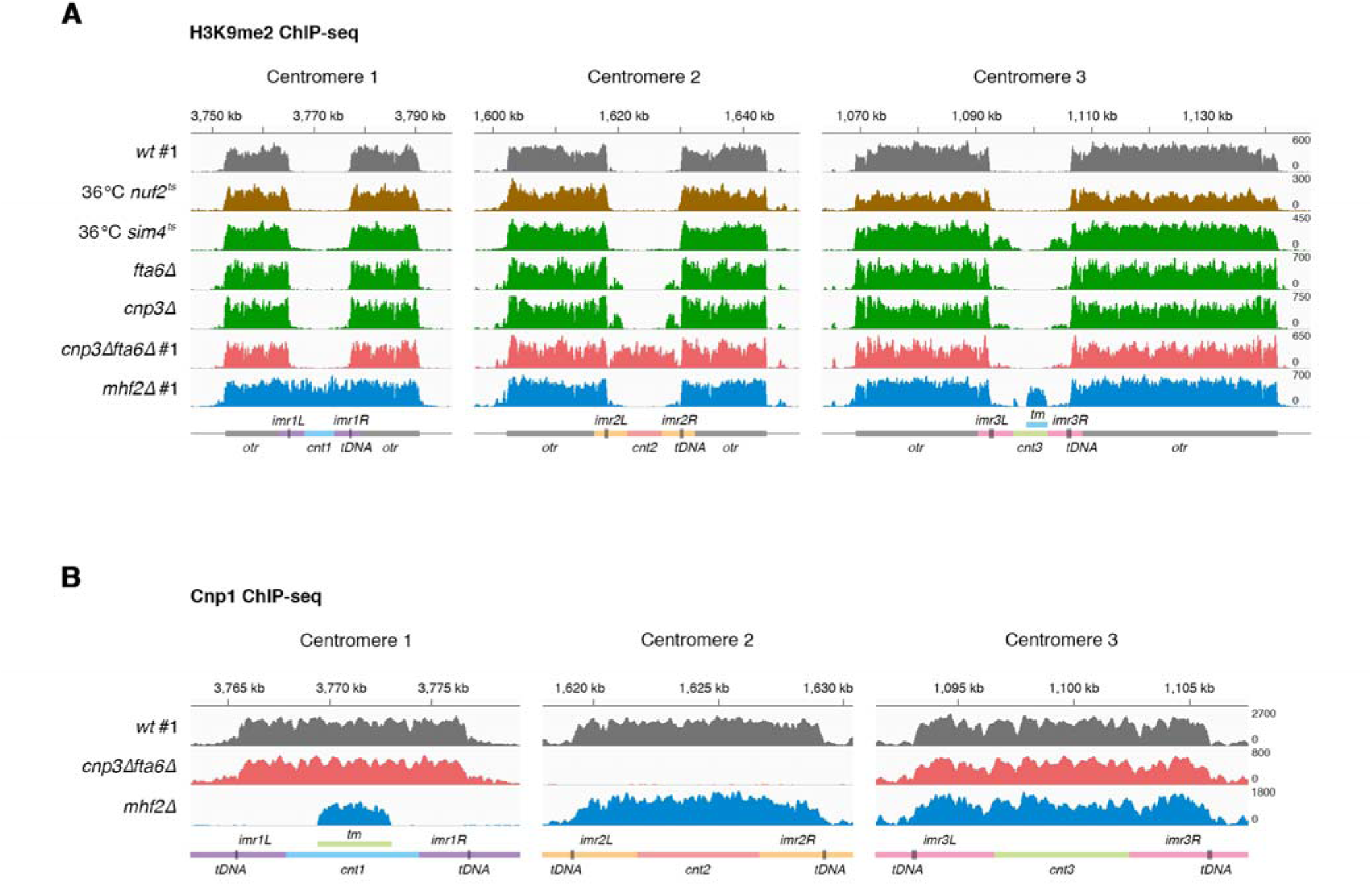
*mhf2*Δ and *cnp3*Δ*fta6*Δ induce centromere inactivation. **(A)** H3K9me2 ChIP-seq reads mapped to centromeric and pericentromeric regions of all three chromosomes in outer kinetochore mutants (brown), inner kinetochore mutants (green), *mhf2*Δ (blue) and *cnp3*Δ*fta6*Δ (pink) compared to wild-type cells (gray). Strain names are as labeled (#1, biological replicate 1). *mhf2*Δ (blue) and *cnp3*Δ*fta6*Δ (pink) showed complete occupancy of H3K9me2 on *cnt1* and *cnt2*, respectively. 36 °C, temperature sensitive (*ts*) strains were incubated at 26 °C and shifted to 36 °C for 6 hours. **(B)** Cnp1 ChIP-seq reads mapped to centromeric regions of all three chromosomes in *mhf2*Δ (blue) and *cnp3*Δ*fta6*Δ (pink) compared to wild-type cells (gray). #1, biological replicate 1. Diagrams illustrate the organization of centromere 1, 2 and 3. *tDNA*, vertical lines; *tm*, segments with identical sequences in *cnt1* and *cnt3*. X axis, DNA coordinates on chromosome 1, 2 and 3 according to reference genome (pombase.org); Y axis, reads per million of ChIP-seq reads randomly assigned to the repetitive DNA sequences. The wild-type ChIP-seq raw data were previously published (30).

## Results

### *mhf2*Δ and *cnp3*Δ*fta6*Δ induce centromere inactivation

Here we investigated the influence of kinetochore mutations on centromeric chromatin organization. Anti-H3K9me2 chromatin immunoprecipitation and high-throughput sequencing (ChIP-seq) were performed in mutants with either deletions of non-essential kinetochore genes or conditional inactivation (temperature sensitive, *ts*) mutations in essential ones. We detected pericentromeric heterochromatin spreading into the core regions to various degrees: the outer kinetochore mutants (*nuf2-1* and *mis12-537*) showed no heterochromatin spreading; whereas the inner kinetochore mutants (*mis15-68, sim4-193, mal2-1, fta6*Δ and *cnp3*Δ) exhibited minor to prominent heterochromatin encroaching, and *mhf2*Δ (mammalian CENP-X homolog) exhibited complete heterochromatin occupancy in one centromere (*cen1*) but normal pericentromeric distribution in *cen2* and *cen3* (Fig. 1A and Fig. S1A). We also constructed a double-mutant *cnp3*Δ*fta6*Δ and found its *cen2* completely covered by heterochromatin using anti-H3K9me2 ChIP-seq (Fig. 1A and Fig. S1A), suggesting that perturbations to centromeric chromatin by *cnp3*Δ and *fta6*Δ cumulatively led to centromere inactivation. Anti-Cnp1 ChIP-seq detected no significant Cnp1 signal at the centromeric cores occupied by heterochromatin but wild-type levels of Cnp1 at the other two in both *mhf2*Δ and *cnp3*Δ*fta6*Δ (Fig. 1B), confirming that only *cen1* or *cen2* was inactivated (designated as *cen1*^inactive^ and *cen2*^inactive^ hereafter) in these two strains, respectively. We also examined mutants of genes encoding centromere-interacting proteins (*mis16-53, mis18-262, ams2*Δ and *sim3*Δ) shown to affect centromeric Cnp1 incorporation and detected no noticeable heterochromatin spreading (Fig. S1B) (13-15). These results together demonstrate that the integrity of the inner kinetochore is required to maintain normal centromeric chromatin organization as well as distinct centromere identity.

### Neocentromeres are formed preferably on pericentromeric regions in single depletion of CENP-T-W-S-X

CENP-T-W-S-X is a conserved inner kinetochore complex in which each subunit contains a histone fold domain that binds directly to DNA (16). To further explore the role of the CENP-T-W-S-X complex in maintaining centromere identity, we generated heterozygous deletion diploid strains for the three non-essential components *wip1*/CENP-W, *mhf1*/CENP-S and *mhf2*/CENP-X in *pombe* (*cnp20*/CENP-T is essential) (9). Tetrad analysis of the meiotic progeny of each strain showed that most of the asci contained two or fewer viable spores, demonstrating a significant reduction in meiotic progeny viability (Fig. S2). However, the lethality was not linked to gene deletions among the progeny (Table S1). By anti-Cnp1 ChIP-seq, random inactivation in only one of the three centromeres was found in each of the ten tested *wip1*Δ, *mhf1*Δ or *mhf2*Δ haploid strains (Fig. 2A, Fig. S3A and Table S2), whereas the wild-type progeny from the same asci exhibited normal pericentromeric heterochromatin distribution (Fig. 2B and Fig. S3B). No strain was found carrying more than one inactivated centromere. We speculate that simultaneous inactivation of two or three centromeres may not be tolerated for cell survival. Normal pericentromeric heterochromatin distribution was also detected in the parental heterozygous diploid cells (Fig. 2B), excluding the possibility that centromere inactivation occurred pre-meiotically. In rare (6.1%) *mhf2*Δ/*+* asci containing four viable spores, two *mhf2*Δ progeny carrying different inactivated centromeres (*cen2*^inactive^ and *cen3*^inactive^, respectively) were recovered from the same ascus, further supporting the notion that centromere inactivation occurred independently and postzygotically (Fig. S3C).

**Fig. 2.**
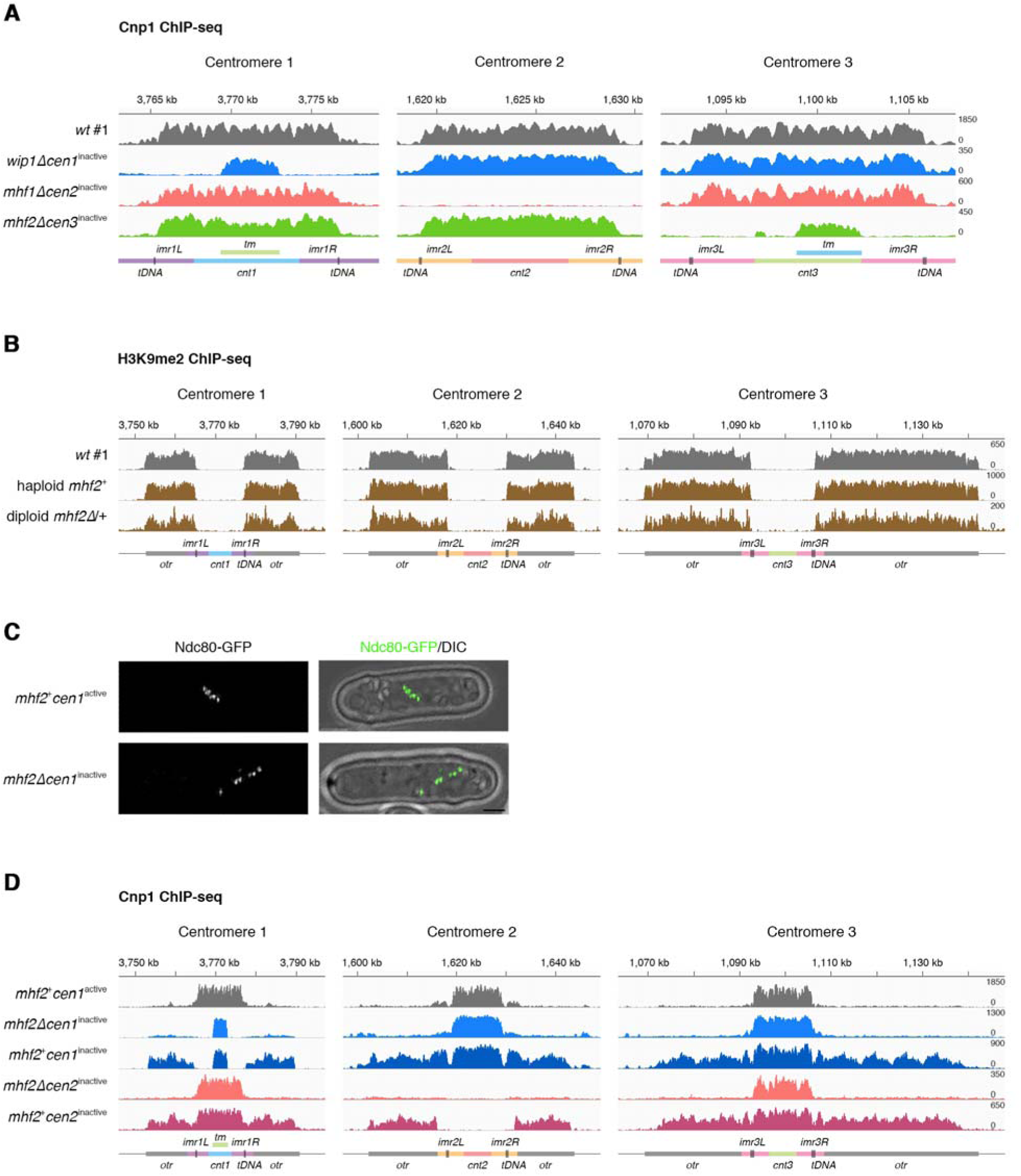
Neocentromeres are formed preferably on pericentromeric regions in single depletion of CENP-T-W-S-X. **(A)** Cnp1 ChIP-seq reads mapped to centromeric regions of all three chromosomes in randomly chosen *wip1*Δ, *mhf1*Δ and *mhf2*Δ strains (*cen1*^inactive^ blue, *cen2*^inactive^ pink, *cen3*^inactive^ green) compared to wild-type strain (*cen1*/*2*/*3*^active^ gray). **(B)** H3K9me2 ChIP-seq reads mapped to centromeric and pericentromeric regions of all three chromosomes in meiotic haploid progeny *mhf2*^+^ and heterozygous deletion diploid *mhf2*Δ/+ (brown) compared to wild-type cells (gray). **(C)** Six dots of outer kinetochore Ndc80-GFP observed in *mhf2*+*cen1*^active^ (upper panels) and *mhf2*Δ*cen1*^inactive^ (lower panels) M phase cells treated with the thiabendazole (TBZ, 20 μg/ml). Scale bar, 2 μ*m*. **(D)** Cnp1 ChIP-seq reads mapped to centromeric and pericentromeric regions of all three chromosomes in *mhf2*Δ*cen1*^inactive^ (blue) and *mhf2*^+^*cen1*^inactive^ (dark blue); *mhf2*Δ *cen2*^inactive^ (pink) and *mhf2*^+^*cen2*^inactive^ (dark pink) compared to wild-type cells (gray). Diagrams, X axis and Y axis, same as **Fig. 1**.

A previous study in fission yeast has shown that with complete excision of *cen1* DNA by genome editing, a few cells (about 0.5%) survive by either forming a neocentromere at a new location or fusing the acentric chromosome to another chromosome (17). To determine whether the surviving *mhf2*Δ cells acquired a neocentromere, we microscopically examined *mhf2*Δ*cen1*^inacitve^ cells expressing a green fluorescent protein-tagged outer kinetochore protein Ndc80 (Ndc80-GFP) for the presence of a complete set of kinetochores (three pairs of sisters). Six discrete dots were resolved in the M phase cells with their kinetochores scattered sufficiently (Fig. 2C), suggesting that a functional kinetochore (and presumably a neocentromere) was formed on chromosome 1 carrying *cen1*^inacitve^.

To determine the locations of the neocentromeres, anti-Cnp1 ChIP-seq was performed in *mhf2*Δ*cen1*^inactive^ and *mhf2*Δ*cen2*^inactive^. While normal levels of Cnp1 were present in the two functional centromeres, modest but clearly detectable Cnp1 appeared in pericentromeric regions of the inactivated centromere (Fig. 2D). These neocentromeres are very likely to be functional considering that kinetochores successfully formed on all chromosomes (Fig. 2C). The Cnp1 signals from ChIP-seq indicate that either Cnp1 incorporation at the neocentromeres is low in *mhf2*Δ or the positions of Cnp1 nucleosomes might be divergent among individual *mhf2*Δ cells within a population.

### Incompatibility between neocentromeres and endogenous centromeres causes a meiosis barrier

By crossing *mhf2*Δ*cen1*^inactive^ to *mhf2*+ *cen1*^active^, we recovered *mhf2*+ *cen1*^inactive^ among the progeny (Table 1 and Fig. S3D). This suggests that the inactivated centromere (and presumably the accompanied neocentromere) can be inherited even in the absence of the genetic lesion that induced it. In *mhf2* + *cen1*^inactive^ and *mhf2* + *cen2*^inactive^, Cnp1 signals comparable to that at the endogenous centromeres, were detected in pericentromeric regions of the inactivated centromeres by anti-Cnp1 ChIP-Seq (Fig. 2D). Together, these results demonstrated that pericentromeric heterochromatin was the most preferable site for neocentromere seeding. This is in sharp contrast to the scenario in which the neocentromere was formed near the subtelomeric heterochromatin regions upon complete removal of *cen* DNA including pericentromeric repeats (17). Due to high DNA sequence similarity between the pericentromeric repeats (18), we were unable to distinguish whether neocentromeres were formed only on one side or symmetrically on both sides of the centromeric cores, or whether Cnp1 occupancy on the *otr* repeats is limited to one centromere or ubiquitous for all three centromeres.

**Table 1.** Incompatibility between neocentromeres and endogenous centromeres causes a meiosis barrier. Cells with heterozygous (*mhf2*Δ*cen1*^inactive^ × *mhf2*+*cen1*^active^, *mhf2*+*cen1*^inactive^ × *mhf2*+*cen1*^active^, *mhf2*+*cen1*^inactive^ × *mhf2*+ *cen2*^inactive^) or homozygous (*mhf2*+ *cen1*^active^ × *mhf2*+ *cen1*^active^, *mhf2*+ *cen1*^inactive^ × *mhf2*+ *cen1*^inactive^) centromeres were crossed and subjected to tetrad dissection. Intact asci with four spores were dissected microscopically and scored for the number of viable spores. Details are listed in Table S4. Spore viability is calculated as the ratio of the number of viable spores to the number of analyzed spores. > 50% reduction in spore viability is defined as meiosis barrier.

| Cross | Spore viability | Dissected<br>Asci (n) | Asci with 4<br>viable spores | Asci with one or<br>no viable spore | Meiosis<br>barrier |
| --- | --- | --- | --- | --- | --- |
| $mhf2^+ cen1^{active} \times mhf2^+ cen1^{active}$ | 93.1% | 130 | 80% | 1.54% | × |
| $mhf2\Delta cen1^{inactive} \times mhf2^+ cen1^{active}$ | 36.5% | 254 | 5.9% | 53.1% | √ |
| $mhf2^+ cen1^{inactive} \times mhf2^+ cen1^{active}$ | 25.2% | 272 | 4.04% | 69.5% | √ |
| $mhf2^+ cen1^{inactive} \times mhf2^+ cen1^{inactive}$ | 89.8% | 108 | 68.5% | 1.9% | × |
| $mhf2^+ cen1^{inactive} \times mhf2^+ cen2^{inactive}$ | 19.8% | 377 | 0.8% | 81.2% | √ |

When crossed to wild type (*cen1*/2/3^active^), both *mhf2*Δ*cen1*^inactive^ and *mhf2*+*cen1*^inactive^ showed high lethality in meiotic progeny. Strikingly, however, homozygotic meiosis (here, crossing between cells carrying the same neocentromere) exhibited near or at wild-type levels of spore viability (Table 1 and Table S3), suggesting that *mhf2*Δ and *cen1*^inactive^ were competent for meiosis. We further performed a series of genetic crosses among strains with different or same centromeres and characterized spore viability (Table 1 and Table S3). Together, the results demonstrated that incompatibility between the neocentromere and the endogenous centromere alone caused poor spore viability. Consistently, microscopic observation of the GFP-labeled chromosome 1 distribution among meiotic progeny demonstrated that severe chromosome segregation defects occurred in both meiosis I and II in diploids carrying mismatched centromeres (Fig. S4 A and B). The specific cause(s) for these meiosis defects remains unclear and may be due to compromised processes related to homologous centromeres such as centromere pairing in early meiosis, or compromised recombination repression at centromeres (19). Overall, mismatching in centromere position between the homologous chromosome pair(s) causes hybrid infertility and constitutes a meiosis barrier between the two strains.

### The endogenous centromere 1 in *mhf2*^+^ is converted to the inactivated centromere 1 by *mhf2*Δ through genetic crossing

In the genetic crosses above, a few asci produced four viable progeny, allowing reliable analysis of the inheritance of genetic lesions and epigenetic features. As expected, *mhf2*Δ conformed to Mendelian inheritance. Likewise, in *mhf2*+*cen1*^inactive^ × *mhf2*+*cen1*^active^, *cen1*^inactive^ also conformed to Mendelian inheritance, suggesting that *cen1*^inactive^ was meiotically stable (Fig. 3A and Fig. S5A). In contrast, in *mhf2*Δ*cen1*^inactive^ × *mhf2*+*cen1*^active^, all viable progeny tested (8 haploid progeny of two intact tetrads plus 3 haploid progeny of incomplete tetrads) contained *cen1*^inactive^, regardless of whether *mhf2*+ or *mhf2*Δ was in the haploid genome (Fig. 3B and Fig. S5 B and C). Thus, in this genetic cross, *cen1*^active^ had a high propensity to be converted into *cen1*^inactive^, most likely using *cen1*^inactive^ on the homologous chromatin as the template. Although the mechanism remains unclear, this centromere conversion phenomenon underscores the pivotal role of *mhf2* in maintaining centromere identity and the consequence of *mhf2*Δ to facilitate the propagation of the neocentromere in the cell population.

**Fig. 3.**
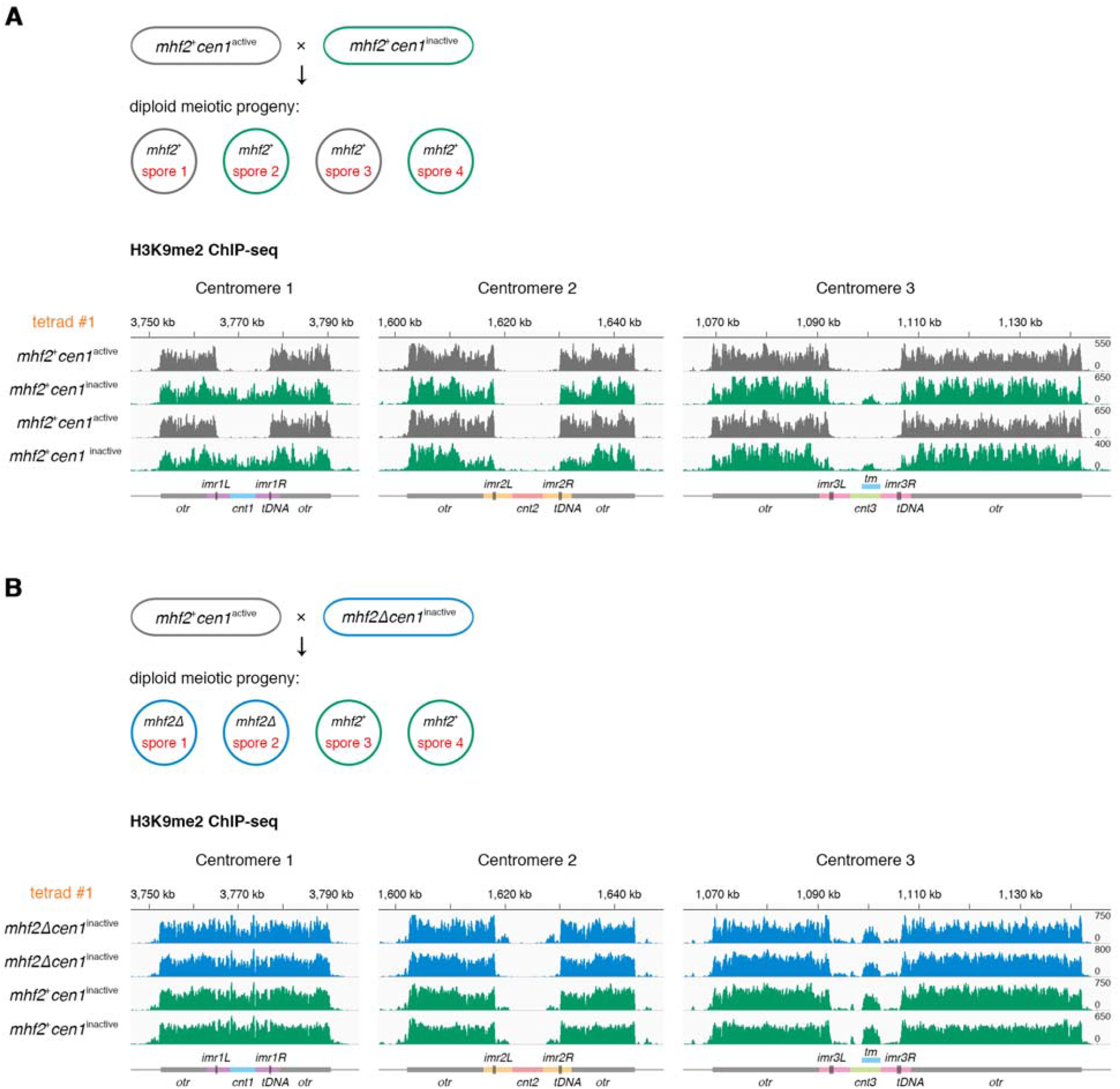
The endogenous centromere 1 in *mhf2*^+^ is converted to the inactivated centromere 1 by *mhf2* Δ through genetic crossing. **(A)** Diagram illustrating the meiotic progeny of *mhf2*+*cen1*^inactive^ × *mhf2*+*cen1*^active^ in asci with four viable spores. H3K9me2 ChIP-seq reads mapped to centromeric and pericentromeric regions of all three chromosomes in four viable progeny from the same ascus (tetrad #1). *cen1*^inactive^ (green) conformed to Mendelian inheritance (2: 2 segregation pattern). **(B)** Same procedure as **(A)** for analyzing *mhf2*Δ*cen1*^inactive^ × *mhf2*+*cen1*^active^. *mhf2*Δ conformed to Mendelian inheritance (2: 2 segregation pattern). Only *cen1*^inactive^ was detected in *mhf2*Δ (blue) and *mhf2*+ (green). Diagrams, X axis and Y axis, same as **Fig. 1.**

We further investigated whether heterochromatin spreading had a causal effect on centromere inactivation by exploring the impact of deletion of *clr4,* which encodes the only heterochromatin modification enzyme (H3K9 methyltransferase) in *pombe* (5). The results suggested that induction of centromere inactivation does not require heterochromatin (Fig. S6). Thus, instead of being a causal factor, heterochromatin spreading is likely the consequence of centromere perturbation and inactivation. It is possible that other molecular features or processes in the pericentromeric regions such as ncRNA transcription or siRNA processing may favor neocentromere formation.

## Discussion

Centromere positioning is remarkably stable in all eukaryotes although neocentromere formation due to centromere repositioning without the incurring of chromosomal marker order variation does occur sporadically in contemporary populations (*e.g.*, about 100 cases of neocentromeres have been reported in humans) (20) and among related species in the evolutionary time scale (*i.e.*, Evolutionary New Centromeres, ENCs) (21). Our study shows that genetic abrogation of the fission yeast CENP-T-W-S-X complex readily initiates centromere repositioning and facilitates the propagation of neocentromeres in the population through meiosis but is dispensable for the mitotic maintenance of neocentromeres (Fig. S7 A and B). A recent study also reported that in cultured chicken DT40 cells, knocking out non-essential constitutive kinetochore components including CENP-S enhances centromere drift upon prolonged cell proliferation (22). Together, these results reveal a fundamental and evolutionarily conserved role of the kinetochore in maintaining centromere identity, and suggest an efficient means of inducing neocentromere formation without incurring centromeric DNA changes, or chromosomal rearrangements (23). The fact that relatively few neocentromeres have been detected so far may be explained by experimental limitations in detection and/or their elimination due to possible detrimental effects on cell fitness. Functionally, mismatching between the original centromere and the neocentromere alone is sufficient to impose a meiosis barrier between the two populations, with an efficiency comparable to other known mechanisms in fission yeast including chromosomal rearrangements (24) and spore killer genes (25, 26). We thus propose that neocentromeres seen as ENCs may represent an initiation step for genetic divergence during speciation (27-29).

## Acknowledgements

We thank Robin Allshire for providing the Cnp1 antibody. This work was supported by National 973 Plan for Basic Research Grant 2015CB910602 (to X. H.) and National Natural Science Foundation of China (NSFC) Grant 31628012 (to X. H.).

## Author contribution

M.L. conceived the project and performed the experiments. M.L. and X.H. designed the study and analyzed the data. M.L. and X.H. wrote the manuscript.

## Competing interests

The authors declare no completing interests.

## List of Supplementary Information

Materials and Methods

Figures S1-S7

Table S1-S5

